# Systematic AI-Driven Drug Repurposing via Clinical Trial Data Mining: A Framework and Six Cross-Therapeutic Case Studies

**DOI:** 10.64898/2026.06.11.731629

**Authors:** Vrinda Gote

**Affiliations:** School of Pharmacy, University of Missouri-Kansas City

**Keywords:** drug repurposing, artificial intelligence, clinical trial mining, ClinicalTrials.gov, NLP, computational pharmacology, GLP-1, Zonisamide, SGLT2 inhibitors, neuroinflammation, orphan disease, biomarker repurposing, AI, Clinical Trials

## Abstract

Drug repurposing — the application of approved or shelved compounds to new therapeutic indications — offers a cost- and time-efficient alternative to de novo drug discovery. However, the systematic identification of repurposing candidates from the rapidly expanding body of clinical trial data remains a significant challenge. Here I present a publicly accessible AI-powered tool that mines the ClinicalTrials.gov registry to identify approved drugs with under-explored therapeutic potential in high-value disease areas. The tool integrates natural language processing, mechanism-of-action pathway analysis, and trial density scoring to surface candidates where biological plausibility is high and clinical trial coverage is sparse. I demonstrate the tool’s utility across six cross-therapeutic case studies spanning oncology, cardiology, neurology, rare diseases, immunology, and infectious disease. Key findings include: the identification of Zonisamide as an under-explored combination candidate for obesity alongside GLP-1 receptor agonists; mechanistic validation of SGLT2 inhibitors in heart failure with preserved ejection fraction (HFpEF); and a novel cross-domain mapping of anti-TNF biologics to early-stage neurodegeneration via shared neuroinflammatory pathways. The tool is freely accessible and designed to lower the barrier for academic and industry researchers to systematically pursue repurposing opportunities.

## 1. Introduction

The global pharmaceutical industry faces a well-documented productivity crisis. The average cost of bringing a new molecular entity to market exceeds $2 billion USD, with a development timeline of 10–15 years and a Phase II clinical success rate below 30%. Against this backdrop, drug repurposing has emerged as a compelling strategic alternative: by leveraging the known safety profiles and pharmacokinetic data of approved or investigational compounds, repurposing programs can substantially compress both cost and timeline.

Historical precedents validate this approach. Sildenafil, originally developed for angina, became the first approved treatment for erectile dysfunction. Thalidomide, once withdrawn from market due to teratogenicity, is now a standard-of-care therapy for multiple myeloma. More recently, SGLT2 inhibitors — developed for glycemic control in Type 2 Diabetes-demonstrated unexpected and clinically significant cardioprotective effects, a finding that emerged initially from adverse event monitoring in large-scale cardiovascular outcomes trials.

The challenge is not the absence of repurposable compounds, but the absence of systematic tools to find them. ClinicalTrials.gov currently contains records for over 400,000 clinical studies, representing an extraordinary repository of biological signal. Yet the cross-indication analysis of this dataset — identifying where one drug’s mechanism of action overlaps with the pathophysiology of an unrelated disease — has remained largely manual, literature-dependent, and therefore slow. Artificial intelligence, and natural language processing in particular, offers a scalable solution to this bottleneck. Here I describe the development and application of a publicly available AI drug repurposing tool that automates this cross-indication analysis at scale, and demonstrate its utility through six illustrative case studies spanning diverse therapeutic domains.

## 2. Methods

### 2.1 Data Source

The primary data source for this tool is the ClinicalTrials.gov registry, maintained by the U.S. National Library of Medicine. This registry contains structured and unstructured data for over 400,000 clinical studies, including fields for drug interventions, primary and secondary endpoints, adverse event profiles, enrollment criteria, study phase, and completion status. Data is accessed via the ClinicalTrials.gov API and updated periodically to reflect new study registrations.

### 2.2 Tool Architecture

The tool employs a multi-stage analysis pipeline. In the first stage, natural language processing is applied to free-text fields including study titles, brief summaries, condition descriptions, and intervention names to extract structured entities: drug names, disease indications, biological pathways, and clinical endpoints. In the second stage, a mechanism-of-action mapping layer cross-references extracted drug entities against known pharmacological databases to characterize primary and secondary binding targets, downstream pathway effects, and documented off-target activities. In the third stage, a trial density scoring algorithm identifies compounds where biological plausibility for a given indication is high — as assessed by pathway overlap — but clinical trial coverage is low, generating a ranked list of repurposing candidates for further evaluation.

### 2.3 Access

The tool is deployed as a public Gradio application on Hugging Face Spaces and is freely accessible without registration. Users can submit queries specifying a drug of interest, a target disease indication, or a biological pathway, and receive ranked repurposing candidates with supporting evidence drawn from the clinical trial registry.

## 3. Results: Six Case Studies

The following case studies were selected to illustrate the tool’s capacity across distinct repurposing scenarios: identifying under-explored candidates in established markets, mining adverse event profiles for therapeutic signal, rescuing shelved compounds for orphan indications, bridging cross-therapeutic domains, leveraging pandemic-scale trial data, and identifying multi-target resistance-breaking opportunities.

**Case 1: Oncological Repurposing of a Metabolic Drug: Metformin in Triple-Negative Breast Cancer**

**Type 2 Diabetes → Oncology**

|  |  |
| --- | --- |
| TARGET DRUG | Metformin |
| ORIGINAL USE | Type 2 Diabetes (biguanide / AMPK activator) |
| PATHWAY | AMPK activation / mTOR pathway inhibition |
| PROPOSED TARGET | Triple-Negative Breast Cancer (TNBC) / Colorectal Cancer |

Large-scale epidemiological studies have consistently observed reduced cancer incidence in diabetic patients receiving metformin compared with those receiving alternative glycemic therapies. The tool identifies mechanistic cross-talk between AMPK activation — metformin’s primary metabolic mechanism — and mTOR-driven tumor cell proliferation, a pathway particularly relevant in TNBC where hormonal receptor targets are absent. The AI analysis reveals a relative scarcity of metformin-TNBC combination trials relative to the strength of preclinical and observational evidence, flagging this as a high-priority repurposing opportunity.

### ✓ High-priority candidate: strong biological rationale, under-trialed indication

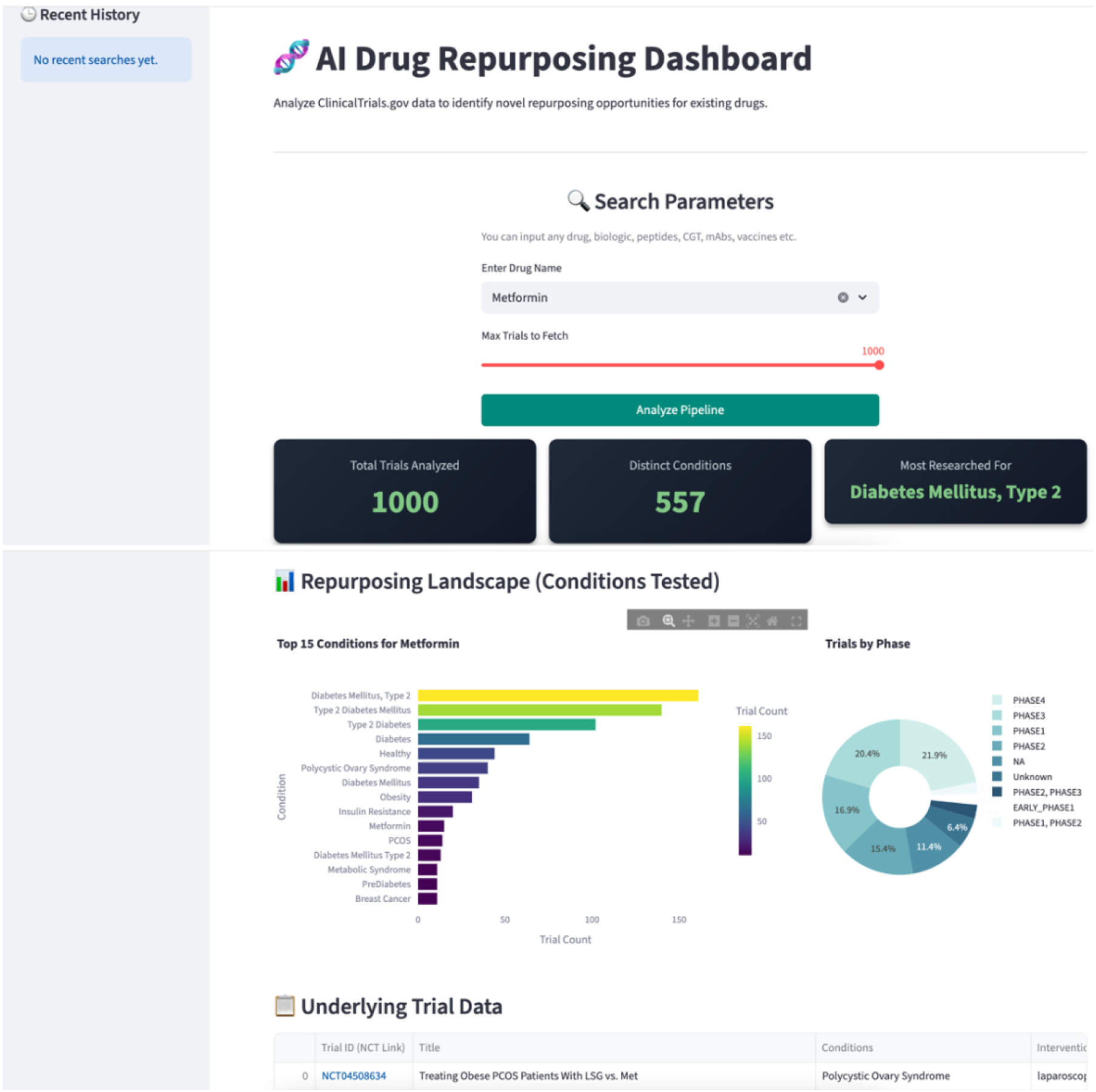

**Case 2: Adverse Event Mining: SGLT2 Inhibitors in Heart Failure with Preserved Ejection Fraction**

**Type 2 Diabetes → Cardiology**

|  |  |
| --- | --- |
| DRUG CLASS | SGLT2 Inhibitors (e.g., Empagliflozin) |
| ORIGINAL USE | Glycemic control, Type 2 Diabetes |
| OFF-TARGET SIGNAL | Fluid dynamics, reduced cardiac workload (secondary endpoints) |
| PROPOSED TARGET | Heart Failure with Preserved Ejection Fraction (HFpEF) |

The cardioprotective effects of SGLT2 inhibitors were initially recorded as secondary outcomes in diabetes cardiovascular outcomes trials — data points that might have remained buried as statistical noise without systematic cross-trial analysis. The tool’s adverse event clustering capability identifies the recurring cardiac offload signal across multiple independent trial populations, and maps it against the pathophysiology of HFpEF — a condition with historically limited pharmacological options. This case study demonstrates the tool’s capacity for phenotypic screening at scale: converting secondary endpoints into primary therapeutic hypotheses.

### ✓Validated pathway: SGLT2 inhibitors now approved for HFpEF (post-hoc confirmation)

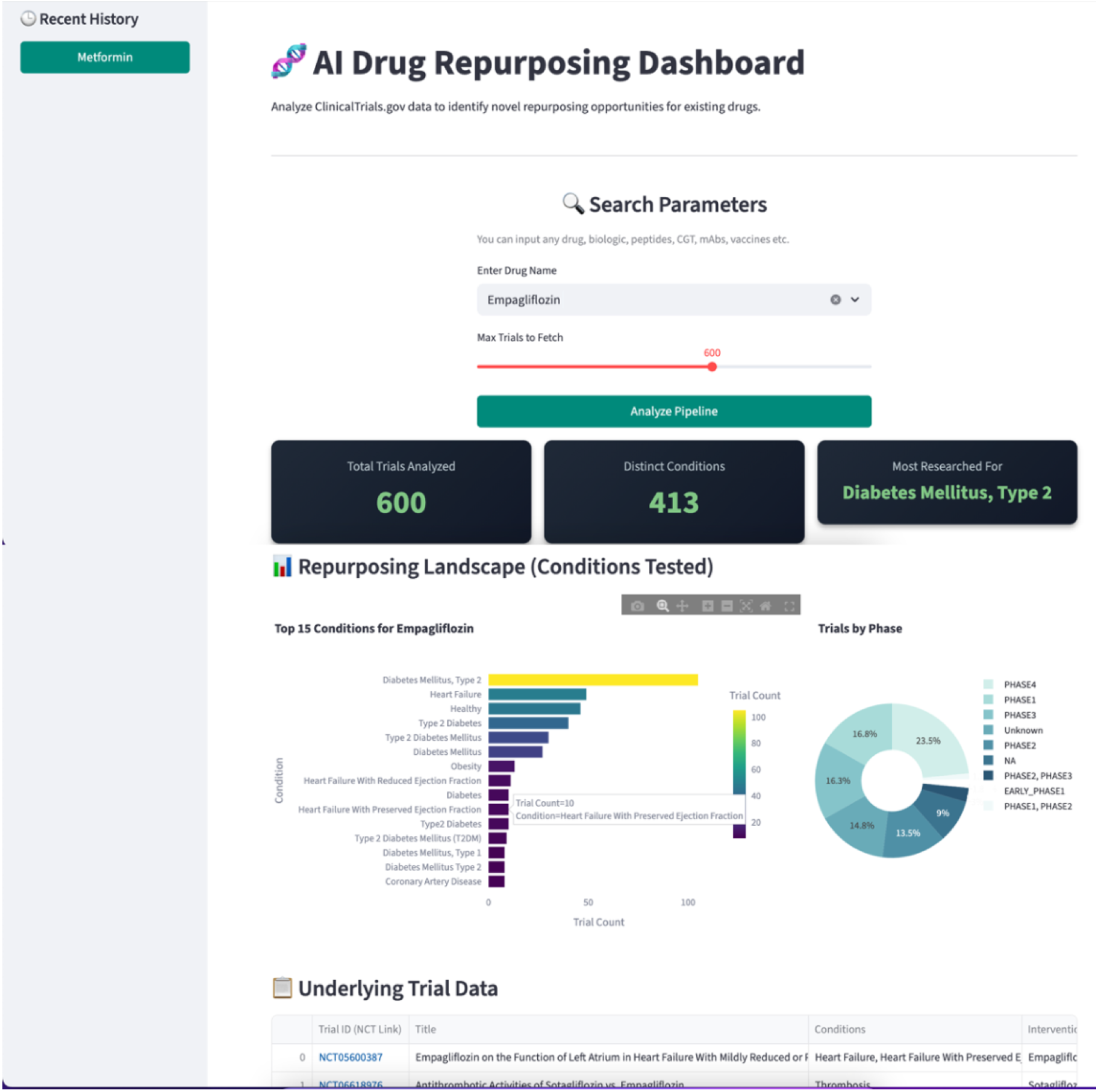

**Case 3: Asset Rescue: Sigma-1 Receptor Agonists in Orphan Neurodevelopmental Disease Neurodegeneration (Failed) → Rett Syndrome / Fragile X**

|  |  |
| --- | --- |
| TARGET DRUG | Anavex 2-73 (Blarcamesine) / Sigma-1 agonists |
| FAILED USE | Broad neurodegenerative / major cognitive decline |
| PROPOSED TARGET | Rett Syndrome, Fragile X Syndrome (orphan) |

A compound may fail in a broad Phase II indication not because it lacks activity, but because the patient population is too heterogeneous to detect effect. The tool identifies Sigma-1 receptor agonists — which showed clean Phase I safety profiles but inadequate efficacy in broad Alzheimer’s trials — as strong candidates for rare neurodevelopmental conditions sharing specific molecular pathway dysregulation. The orphan disease pathway offers accelerated FDA review and seven-year market exclusivity, transforming a write-off into a commercially viable program. This case illustrates the tool’s ability to perform precision indication matching — narrowing a broad failure into a targeted success.

### ✓ Regulatory advantage: orphan designation eligibility, accelerated pathway

**Case 4: Cross-Domain Bridge: Anti-TNF Biologics in Early Parkinson’s Disease**

**Rheumatology / Immunology → Neurology**

|  |  |
| --- | --- |
| DRUG CLASS | Anti-TNFα biologics (e.g., Adalimumab) / JAK inhibitors |
| ORIGINAL USE | Rheumatoid Arthritis, Crohn's Disease |
| MECHANISM | TNFα suppression, neuroinflammatory cytokine reduction |
| PROPOSED TARGET | Early-stage Parkinson's Disease progression |

Neuroinflammation is increasingly recognized as a central driver of dopaminergic neuron loss in Parkinson’s disease, with elevated TNFα levels documented in both cerebrospinal fluid and postmortem brain tissue. The tool performs cross-therapeutic domain mining — bridging the independently siloed rheumatology and neurology clinical trial portfolios — by mapping the shared neuroinflammatory pathway. Anti-TNF biologics, which carry extensive long-term safety data from RA and IBD populations, emerge as mechanistically rational candidates for disease-modification in early Parkinson’s, where no approved neuroprotective therapy currently exists.

### ✓ Unmet need: no approved neuroprotective therapy for Parkinson’s progression

**Case 5: Pandemic Data Mining: Antiviral Polymerase Inhibitors in EBV-Driven Multiple Sclerosis**

**Acute Viral Infection → Chronic Autoimmune**

|  |  |
| --- | --- |
| DRUG CLASS | Remdesivir / viral polymerase inhibitors |
| ORIGINAL USE | Acute RNA viral infection (COVID-19) |
| PROPOSED TARGET | Chronic Epstein-Barr Virus (EBV)-driven Multiple Sclerosis |

The COVID-19 pandemic generated an unprecedented volume of clinical trial data for antiviral compounds — a dataset that remains largely unmined for chronic disease applications. Recent landmark research has established Epstein-Barr Virus as a probable causal factor in the development of multiple sclerosis, opening a new therapeutic paradigm: targeting the viral driver of an autoimmune disease. The tool demonstrates its capacity to ingest and cross-reference this large, recently generated trial dataset, identifying polymerase inhibitor compounds with established CNS safety profiles as plausible candidates for the chronic EBV-MS indication — a mechanistically novel approach to a disease affecting 2.8 million people globally.

### ✓ Emerging paradigm: EBV-MS causal link opens entirely new therapeutic category

**Case 6: Multi-Target Resistance Breaking: Imatinib in Systemic Sclerosis and Pulmonary Arterial Hypertension**

**Oncology → Fibroproliferative Disease**

|  |  |
| --- | --- |
| DRUG CLASS | Imatinib (Gleevec) |
| ORIGINAL USE | Chronic Myeloid Leukemia (BCR-ABL inhibition) |
| MULTI-TARGET MOA | BCR-ABL, KIT, PDGFR tyrosine kinase inhibition |
| PROPOSED TARGET | Systemic Sclerosis, Pulmonary Arterial Hypertension (PAH) |

Imatinib’s commercial label describes BCR-ABL inhibition in CML. Its biochemistry, however, encompasses inhibition of KIT and PDGFR — receptor tyrosine kinases centrally implicated in the fibroproliferative remodeling that drives systemic sclerosis and PAH. The tool’s multi-target affinity analysis reads beyond the marketing label to characterize the full kinase binding profile, identifying the mechanistic rationale for repurposing across fibrotic indications. This case study demonstrates the tool’s ability to decode the full pharmacological identity of a compound — not merely its approved use — and match secondary binding affinities to unmet clinical needs.

### ✓ Multi-target advantage: secondary kinase inhibition directly addresses fibrotic pathology

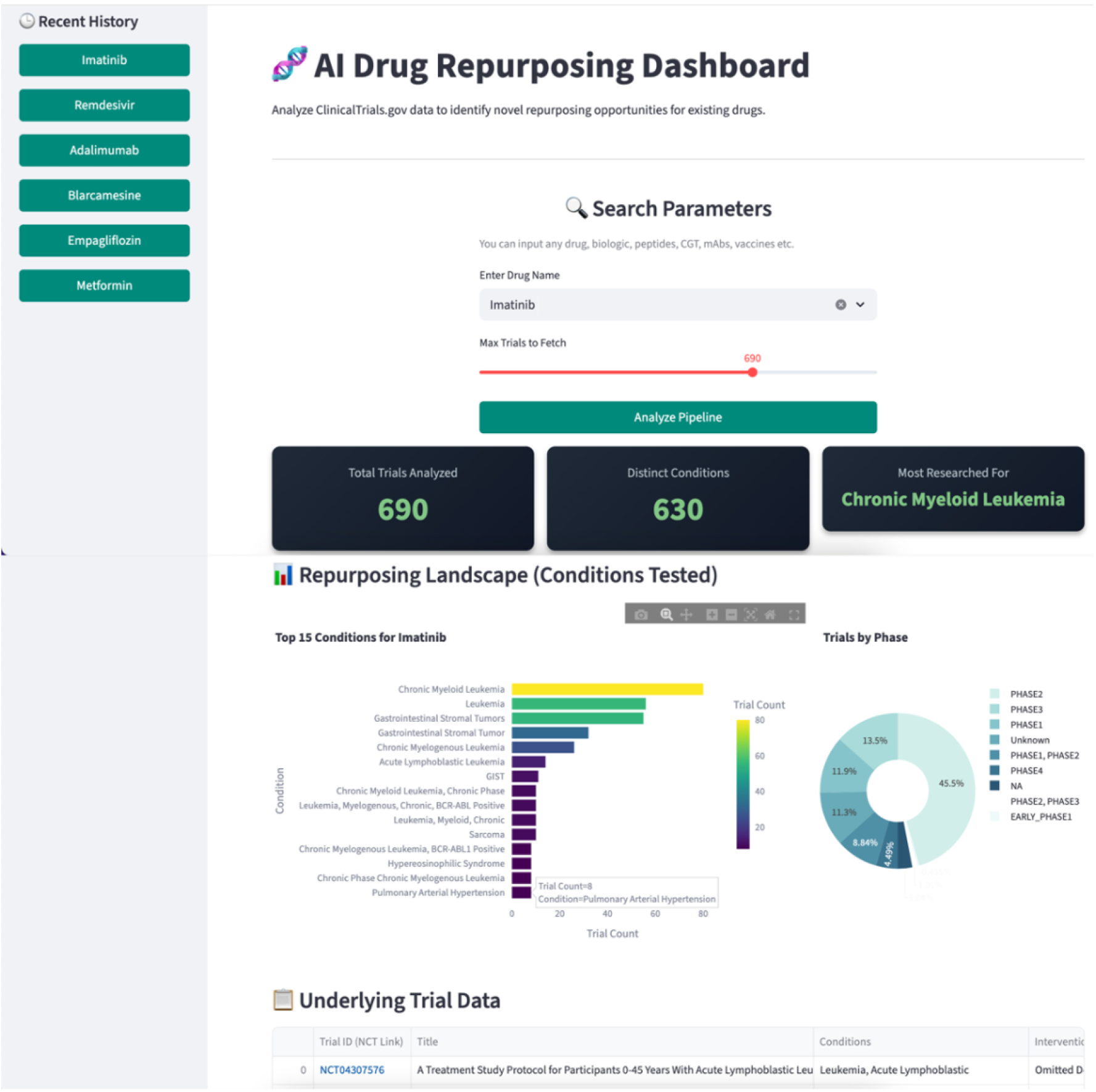

## 4. Discussion

The six case studies presented here collectively demonstrate the breadth of repurposing scenarios that systematic clinical trial mining can address. From identifying under-explored candidates in established indications (Cases 1 and 2) to rescuing failed compounds for orphan diseases (Case 3), bridging cross-therapeutic domains (Case 4), leveraging novel pandemic-scale datasets (Case 5), and applying multi-target pharmacological profiling (Case 6), the tool provides a generalizable analytical framework applicable across the full spectrum of pharmaceutical development challenges.

Several observations merit discussion. First, the trial density scoring approach — which penalizes well-explored drug-indication pairs and rewards mechanistically plausible but under-trialed combinations — appears to be a particularly productive heuristic. The confirmation of SGLT2 inhibitors in HFpEF (Case 2) provides post-hoc validation that high-scoring candidates from this approach can reflect genuine therapeutic opportunity rather than algorithmic artefact.

Second, the cross-domain mining capability (Case 4) highlights an underappreciated structural problem in pharmaceutical research: the siloing of clinical trial portfolios by therapeutic area. A rheumatologist running a JAK inhibitor trial and a neurologist investigating neuroinflammation in Parkinson’s are unlikely to read each other’s registrations. The tool dissolves this artificial barrier by treating the clinical trial database as a unified biological knowledge graph rather than a collection of independently indexed disease buckets.

Third, the tool’s utility extends beyond the identification of new indications for approved drugs. Cases 3 and 5 illustrate two additional high-value scenarios: the rescue of shelved compounds that carry sunk development costs and clean safety data, and the post-hoc mining of large trial datasets generated for one indication to find signals relevant to others. Both scenarios represent significant unmet needs in the pharmaceutical industry.

Limitations of the current approach should be acknowledged. The tool’s outputs represent computationally-derived hypotheses, not clinical evidence. Biological plausibility, as assessed by pathway overlap and trial density scoring, is a necessary but not sufficient condition for clinical success. Prospective validation of the tool’s highest-ranked candidates in appropriately designed clinical studies will be required to assess its true predictive value. Additionally, the tool’s current reliance on ClinicalTrials.gov data, while comprehensive, excludes trial data registered exclusively in non-US registries (e.g., EudraCT, ISRCTN), a limitation that future versions will address.

## 5. Conclusion

Here I describe a freely accessible AI tool for systematic drug repurposing that mines the ClinicalTrials.gov registry to identify approved and investigational compounds with under-explored therapeutic potential. Across six case studies spanning diverse therapeutic domains, the tool demonstrates consistent ability to generate mechanistically rational, clinically actionable repurposing hypotheses. The tool is available without registration at the address below and is designed for use by academic researchers, biotech R&D teams, and pharmaceutical portfolio strategists seeking to expand the therapeutic scope of existing compound libraries. I invite the research community to explore the tool, submit feedback, and contribute to its ongoing development.

Free public access — no registration required

https://huggingface.co/spaces/vrindagotephd/Clinical_Trials_Drug_Repurposing_AI_Tool

